# CARD:Epi – Contextualizing Antimicrobial Resistance Determinants Using Deep Learning Language Models

**DOI:** 10.64898/2026.08.14.744850

**Authors:** Arman Edalatmand, Tiffany E. Ta, Claire Zhao, Abdalmuhaymen Ibrahim, Ramkrishna Upadhyaya, Saduni Rajapaksa, Amogelang R. Raphenya, Andrew G. McArthur

## Abstract

Bacterial outbreak publications outline the key factors involved in the uncontrolled spread of infection. Such factors include the environment, pathogens, hosts, and antimicrobial resistance genes (ARGs). Individually, each paper published in this area gives a glimpse into the devastating impact drug resistant infections have on healthcare, agriculture, and livestock. When examined together, these publications provide contextual information on ARG transmission, from the discovery of new resistance genes to their dissemination to different pathogens, hosts, and environments. We have extracted this information from publications in PubMed by using the biomedical deep-learning language model, BioBERT. We trained BioBERT on two tasks: entity recognition to identify AMR-relevant terms (i.e., ARGs, taxonomy, environments, geographical locations, etc.) and relation extraction to determine which terms identified through entity recognition contextualize ARGs. By collating results from 204,094 antimicrobial resistance publications worldwide, we have generated interpretable results about the sources where genes are commonly found. To visualize the dataset, we have created two pipelines to analyze transmission patterns of ARGs across agriculture, environments, and human populations using a Confusogram and Uniform Manifold Approximation and Projection. Overall, we have taken a large-scale approach to collect antimicrobial resistance data from a commonly overlooked resource, i.e., the systematic examination of the large body of AMR literature and have visualized how scientific literature can be used to assess transmission patterns of ARGs.

## INTRODUCTION

The global rise of antimicrobial resistance (AMR) threatens our ability to successfully treat bacterial infections, undermining modern medicine and contributed to an estimated 1.27 million deaths in 2019.^1^ Despite this, antibiotics are the most commonly prescribed and over-prescribed drugs in human medicine.^1^ The dissemination of antibiotic resistant genes (ARGs) between bacterial communities, whether by a plasmid-mediated, horizontal gene transfer or other mechanism^2,3^ within humans, animals, and environments, contributes significantly to the spread of AMR in a community setting.^4^ Many pathogens (e.g., *Salmonella enterica*, *Escherichia coli*) have complex transmission routes involving multiple environments, necessitating innovative and holistic research solutions.

The One Health model^5,6^ spanning clinical, agricultural, and environmental settings is one such solution which has been adopted by some countries, however, most lack a comprehensive model of ARG transmission dynamics involving farms, livestock, watersheds, food imports, aquaculture, wastewater, wildlife, domestic pets, and human populations. Without these data on transmission, it is challenging to conduct meaningful risk assessments of antibiotics and develop a cohesive national policy for the mitigation of AMR.

Many databases and resources have emerged for structuring genetic and molecular AMR data, such as CARD (the Comprehensive Antibiotic Resistance Database), ResFinder, ARG-ANNOT, and the National Center for Biotechnology Information (NCBI) Pathogen Detection resource.^7–10^ However, while existing genomics tools identify resistance determinants^11^, epidemiological information about these resistance determinants and associated pathogens is beyond their scope. These tools minimally examine the contextual background of resistance determinants, yet public health decisions rely on epidemiological context^12^, such as the geographical distribution and mobility of AMR pathogens and determinants.^13–16^

The most effective method of creating an accurate model of ARG transmission requires genomic epidemiology across the entire One Health spectrum, which has yet to happen; rather, the knowledge of ARG epidemiology is spread piecemeal through scientific literature. Such literature has been growing at a rapid rate; as of 2022, over 33 million publications are stored across PubMed databases, and the number of AMR publications added each year has been growing faster than the PubMed baseline (**Figure 1**).

**Figure 1.**
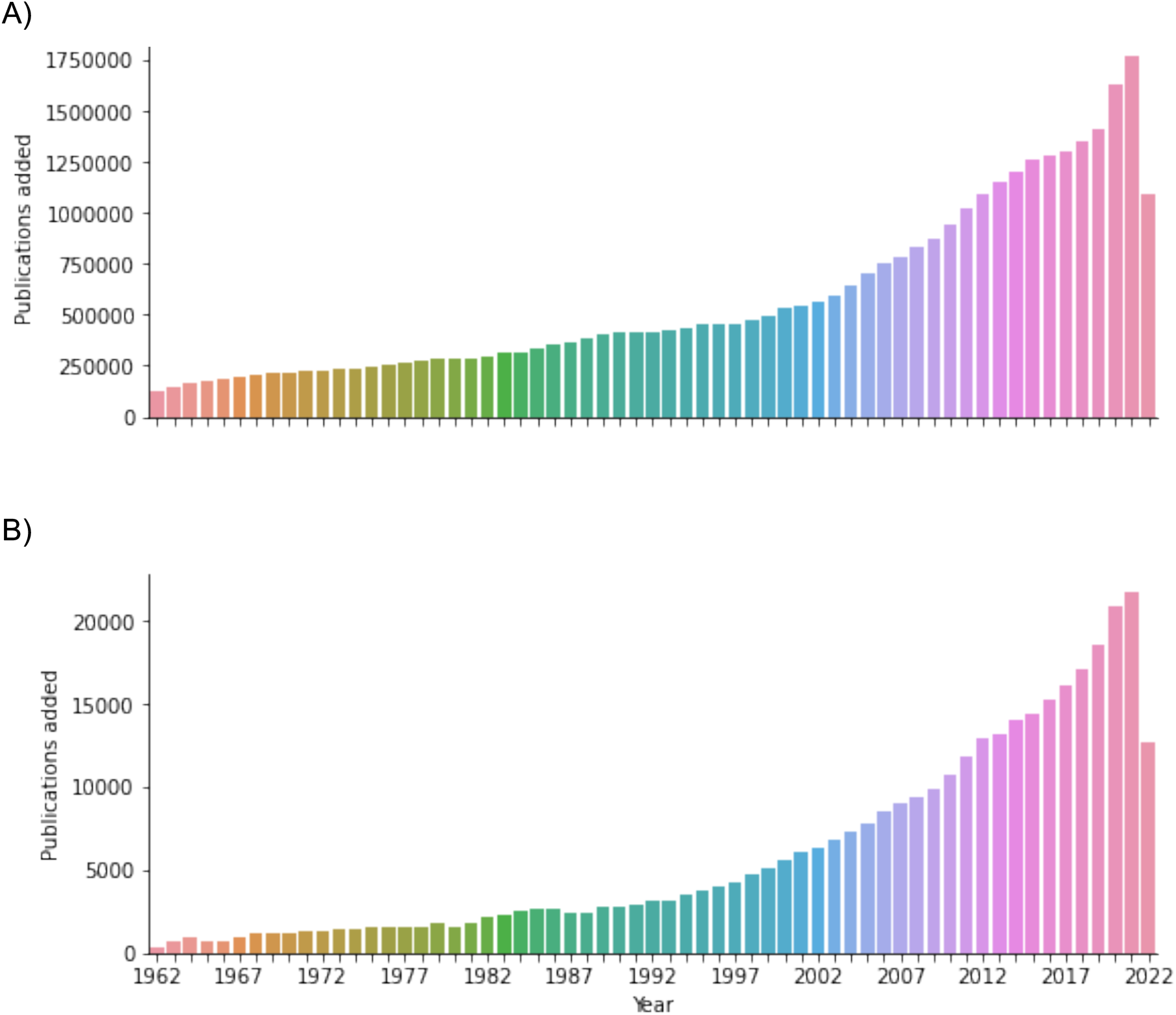
Distribution of publications added to PubMed each year. (A) Total number of publications added to PubMed since 1962, (B) subset of PubMed articles under the PubMed query “antimicrobial resistance”.

Examining the scientific literature may lend sufficient information to model ARG transmission dynamics, however a method is needed to translate disparate literature into data on ARGs. The most efficient way of achieving this is through natural language processing (NLP) to extract words and relationships in biomedical text using named entity recognition (NER) and relation extraction (RE).^17^ The creation of BERT (Bidirectional Encoder Representations from Transformers) solves tasks like NER and RE through a creative method of training.^18^ When dealing with biomedical text, BioBERT can be trained to extract this information from PubMed abstracts.

To extract text, ontologies can be leveraged as they hold a wealth of structuralized information about specific domains of knowledge. The Comprehensive Antibiotic Resistance Database (CARD) is the most cited database of its kind and provides structured information about AMR determinants and antibiotics via the Antibiotic Resistance Ontology (ARO).^19^ To explore the relationships among AMR determinants (from the ARO) and their relationships to epidemiology terms, a diverse set of additional lexicons can be used to extract terms from the literature including the Environment Ontology (ENVO), Food Ontology (FOODON), Gazetteer Ontology (GAZ), Infectious Disease Ontology (IDO), NCBI Taxonomy (NCBI TAXON), Sequence Ontology (SO), and Uber-anatomy ontology (UBERON).^19–27^ The ontological contents can be leveraged for NLP, enabling deep learning to capture useful information.

To collate ARG epidemiological information reported in PubMed, we have created fine-tuned versions of BioBERT that can accurately extract epidemiological and antibiotic resistance determinant terms and relationships from biomedical publications, e.g., gene aph(6)-Id is associated with food-borne pathogens. Additionally, two pipelines using the NER and RE data structures have been designed to visualize these data to assess and predict ARG transmission patterns. This work is a novel extension to CARD, collating AMR-epi biomedical text as CARD:Epi, that allows contextual interpretation of molecular epidemiological information.

## MATERIALS AND METHODS

### Identification of AMR-Epi publications

Using the publicly available PubMed application programming interface (API), we downloaded the abstracts of over 5 million publications from PubMed between 2017 and 2020 to build an AMR-EPI text classifier. To identify publications that contained AMR gene-epidemiological information, we used all publications found under the “Drug Resistance, Bacterial/genetics’ [Mesh]” query in PubMed. Afterwards, they were filtered down to contain publications with a country mention, which would act as the positive set for our model training/testing (3,843 positive publications). For the negative set, random PubMed publication identifiers were generated (10,532 negative publications). 75% of these publications were used for cross-validation, training, and tuning, while the remaining 25% were held out for a final evaluation of the models’ performance.

Publications were preprocessed using the Natural Language Toolkit^28^ to improve model performance. To vectorize text, we used bag-of-words and term frequency-inverse document frequency^29^. Five machine learning models were trained (Logistic Regression, Naïve Bayes, Random Forest, Extreme Gradient Boosting, and Support Vector Machine) with 5-fold cross-validation to prevent overfitting. To further compare each model, an external validation set was created through human curation of a 430-publication subset from September and November 2019. Each curator was given three true/false questions on whether: (1) the publication contained an AMR gene reference, (2) if a geographical location was mentioned in the abstract, and (3) if any other epidemiological information was mentioned. Each publication was assigned two curators; if the two curators disagreed, the publication was excluded.

Once the best performing model and feature extractor pair were identified, the model was tested against the holdout set of 3,594 publications that were not used for model training, cross-validation, or tuning.

### Generating NER and RE training and testing datasets using ontologies

The resource used to gather, find, and download ontologies was the Open Biological and Biomedical Ontology (OBO) Foundry.^30^ Seven ontologies were obtained from the OBO Foundry: ARO^19^, ENVO^21^, FOODON^20^, GAZ, IDO^22^, SO^24^, and UBERON^25^. The lexicon NCBI TAXON^23^ was obtained using the taxonomy database available via the NCBI file-transfer protocol server. **Table S1** displays a detailed breakdown of the lexicons, the concepts that they represent, and the parent term (as a primary filter) that was used.

NER training/testing sets were constructed based on the abstracts of PubMed publications from 2019 passing the AMR-Epi text classifier. Based on the eight lexicons, terms within abstracts were identified using Regular Expressions for text matching. Annotations were filtered based on automated and manual rules. The RE training/testing sets were generated by taking all ARO:other lexicon pair combinations that were found in the same sentence.

### Training BioBERT NER and RE models and normalizing predictions

The NER (8 lexicons) and RE (7 lexicons x ARO) training/testing datasets were used to train BioBERT^31^ models. 25% of data was held out for final evaluation, with the remaining 75% used for training and hyperparameter tuning. Combinations of two hyperparameters, batch size and learning rate, were adjusted to identify the best-performing model, while the epochs remained at a value of 4 (as recommended by the BERT publication).^18^

### Generating the CARD:Epi data set

To generate a full CARD:Epi data set for assessment of possible ARG transmission patterns, the AMR-Epi classifier was run against the abstracts of the 2021 PubMed baseline, i.e., all publications in PubMed with publication date of 2021 or older. This was followed by NER and RE analysis of the positively classified abstracts via our BioBERT models.

To generate interpretable results, annotations gathered from BioBERT NER were mapped to a set of standardized terms. For example, UBERON has multiple terms describing stool, including: “stool”, “feces”, “fecal”, and “faeces”. Matches for these term variants must be mapped to the standard “stool” term. First, Boolean logic was applied to see if the term had an exact match to any terms found within the lexicons. If there was no perfect match, a fuzzy search was conducted by calculating the Levenshtein distance between the annotation and every lexicon term and synonym. An arbitrary cut-off of 95% was used when assigning annotations to lexicon terms. Terms with no perfect match (< 95% cut-off) were passed on to one final fuzzy-matching algorithm. Any terms that were unsuccessful in being normalized were considered “non-normalized” but still included in downstream analyses. For example, the term “broiler houses” was annotated across 56 publications but did not normalize to any term in ENVO.

### Visualization

To visualize the results of RE analyses, gene overlap and Jaccard similarity among terms based on shared ARGs was calculated using custom Python scripts. GraphViz plots were generated using the graphviz library version 0.21 and UMAP plots generated using the umap-learn library version 0.5.11.

## RESULTS

### Identifying drug resistance publications

Various text classifiers (Logistic Regression, Naïve Bayes, Random Forest, Extreme Gradient Boosting, and a Support Vector Machine) were trained and tested with cross-validation to reliably identify publications (via their abstract) containing AMR and epidemiological information. The assumption was that if a publication contained drug resistance and a country name, it would hold additional epidemiological information. Cross-validation resulted in a high receiver operating characteristic (ROC) curve area under the curve of >97% (**Figure 2**). Other indicators of performance (precision, recall, and F1 scores) were also high (>90% F1 score) except for the support vector machine (**Table 1**). The logistic regression model using the term frequency-inverse document frequency (TF-IDF) unigram was selected as the model for all downstream analyses.

**Figure 2:**
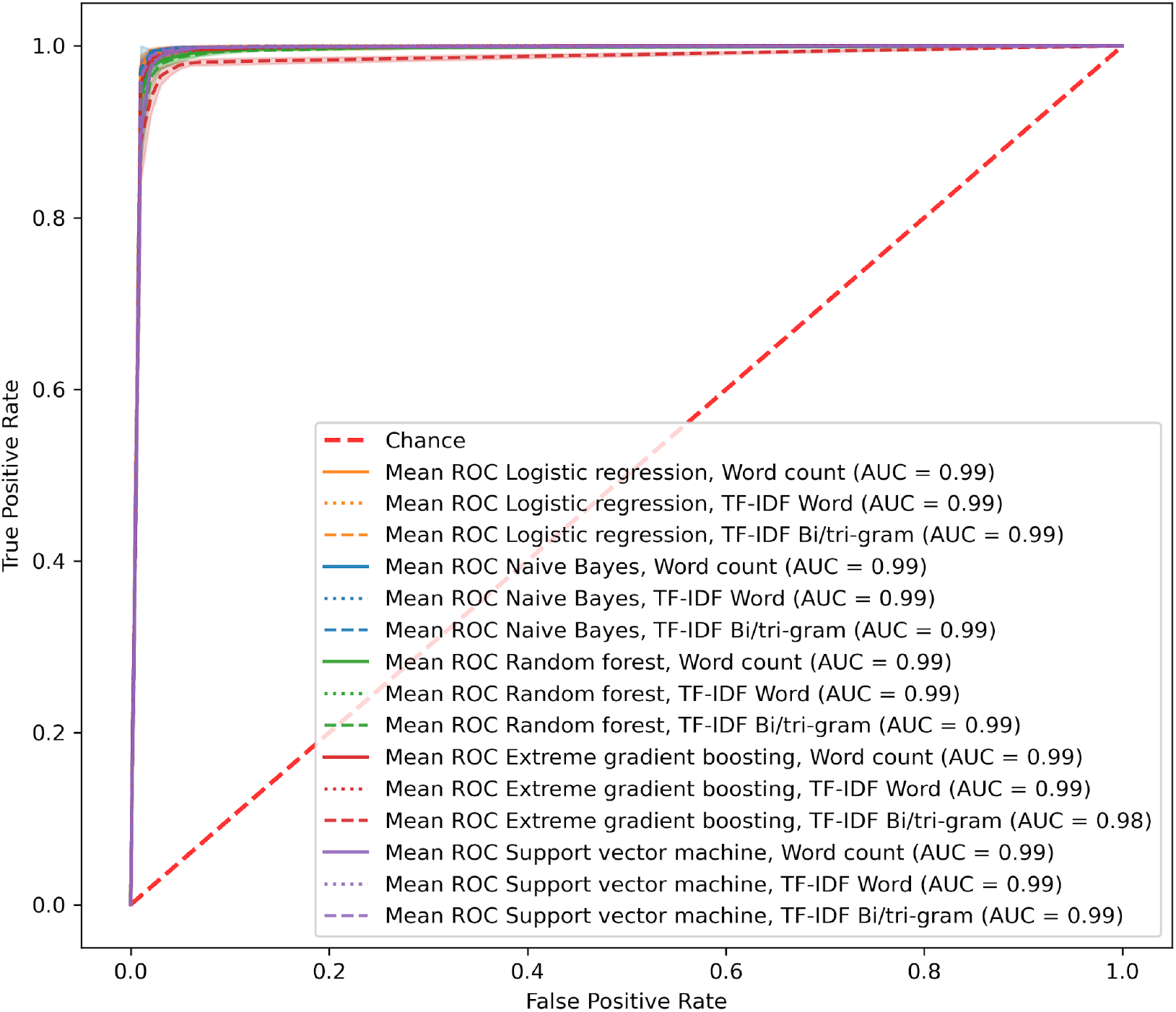
Receiver operator characteristic curve from 5-fold cross-validation of classification models trained on preprocessed abstracts using three feature extraction methods. Results from all five cross-validation tests were averaged to produce a single curve. Shadows around each line are +/- 1 standard deviation from the mean. Models were trained on abstracts that had been preprocessed.

**Table 1:**
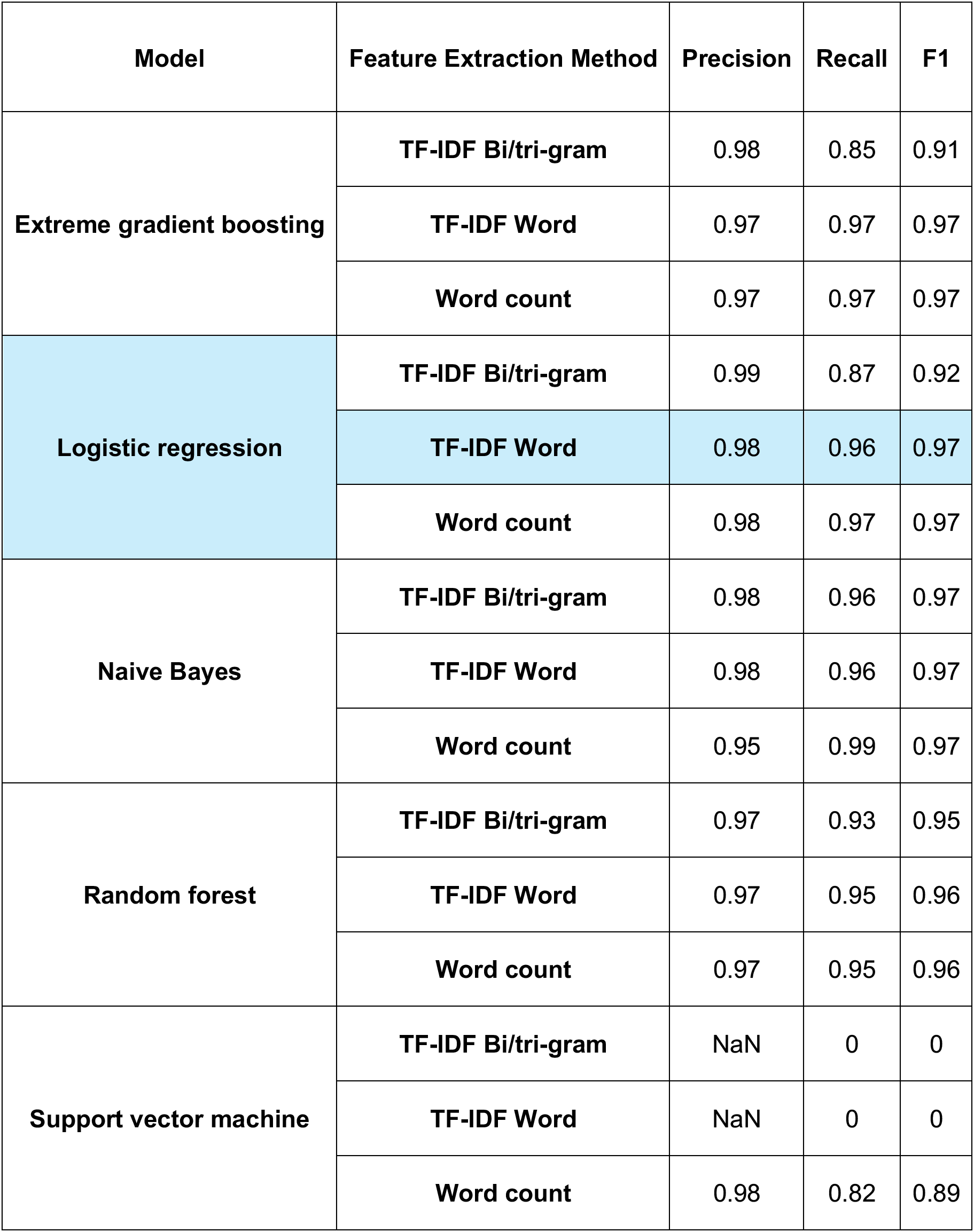
Precision, recall, and F1 performance of AMR-Epi classification models during cross-validation. Highlighted in blue is the final model selected for downstream use.

Since most models performed well according to cross-validation, human curators created an external validation set to additionally assess each model’s performance. When testing the LR TF-IDF Word model on the remaining 25% holdout validation set, it exhibited precision, recall, and F1 values of 99%. As such, it was used to identify all AMR-Epi publications in the 2021 PubMed baseline, yielding 204,094 publications for downstream NER and RE analysis.

### Named Entity Recognition and Relation Extraction deep learning models

To create training and testing datasets for NER, we selected 10,784 publications published in 2019 from the corpus of 204,094 AMR-Epi publications identified. As an initial annotation step, we used regular expressions to match exact lexical terms and their synonyms, to identify 730,538 annotations across 6,839 unique ontology terms (**Table 2**). However, this approach introduced significant error, as many annotations resulted in a mismatch between the annotated term and the corresponding lexicon term’s meaning (e.g., the GAZ term “American Samoa” with the synonym “as”, was erroneously labelled in 10,437 instances of the term where “as” referred to something other than the geographical location). Subsequent manual review revealed the need to include numerous terms that conceptually belonged to a lexicon but were absent from the original ontology and were manually added (e.g., “pork” was identified as relevant but was not present in FOODON). To mitigate these errors, filtering methods were applied and reduced the total number of annotations from 730,538 to 148,445. Based on these curated annotations, eight gold-standard datasets were created, one for each ontology, for fine-tuning BioBERT models to identify both ARGs and epidemiological terms.

**Table 2:** Term, annotation, and publication counts of lexicons in the training data. The term count is the total number of words in a lexicon, not including synonyms. Annotation count is the total number of matches found across all abstracts. The publication count is the number of unique publications in which lexicon terms appear. The number of unique publications with annotations is 10,782 prior to filtering.

| **Lexicon** | **Term count** | **Annotation count** | **Publication count** |
| --- | --- | --- | --- |
| ARO | 3,850 | 10,118 | 2,689 |
| FOODON | 10,815 | 15,529 | 6,682 |
| UBERON | 13,847 | 113,456 | 10,646 |
| ENVO | 217 | 4,866 | 2,633 |
| NCBI TAXON | 466,787 | 43,566 | 8,597 |
| GAZ | 565,454 | 361,759 | 10,780 |
| SO | 2,597 | 146,299 | 10,719 |
| IDO | 274 | 34,945 | 8,793 |

To train a relationship extraction model to identify contextual associations between ARGs and epidemiology terms, we generated 12,588 potential ARG-epidemiology term pairs from our NER training and testing datasets (**Table S2**). These pairs were identified by selecting instances where an ARG and an epidemiology term co-occurred within the same sentence. Subsequently, we manually reviewed each pair to determine if they represented a genuine relationship. This resulted in 10,329 (82%) confirmed relationships and 2,254 (18%) pairs deemed to have no relationships. In total, seven gold-standard datasets were generated for fine-tuning BioBERT models to identify ARG-epidemiological relationships.

After training and testing the NER models, the best performing NER model was determined. A well-performing NER model should generalize predictions, accurately identifying terms belonging to a lexicon even when those terms were not encountered during training. Models trained on lexicons with very few unique terms (ENVO, IDO, SO, and UBERON) demonstrated lower recall for unseen terms (<60%). In contrast, models trained on lexicons with more unique terms, like ARO, FOODON, GAZ, and NCBI TAXON, achieved recall values greater than 70% (**Table 3**). Based on the highest generalization recall value, the best NER model for each lexicon was selected for subsequent downstream predictions (highlighted in **Table 3**).

**Table 3:**
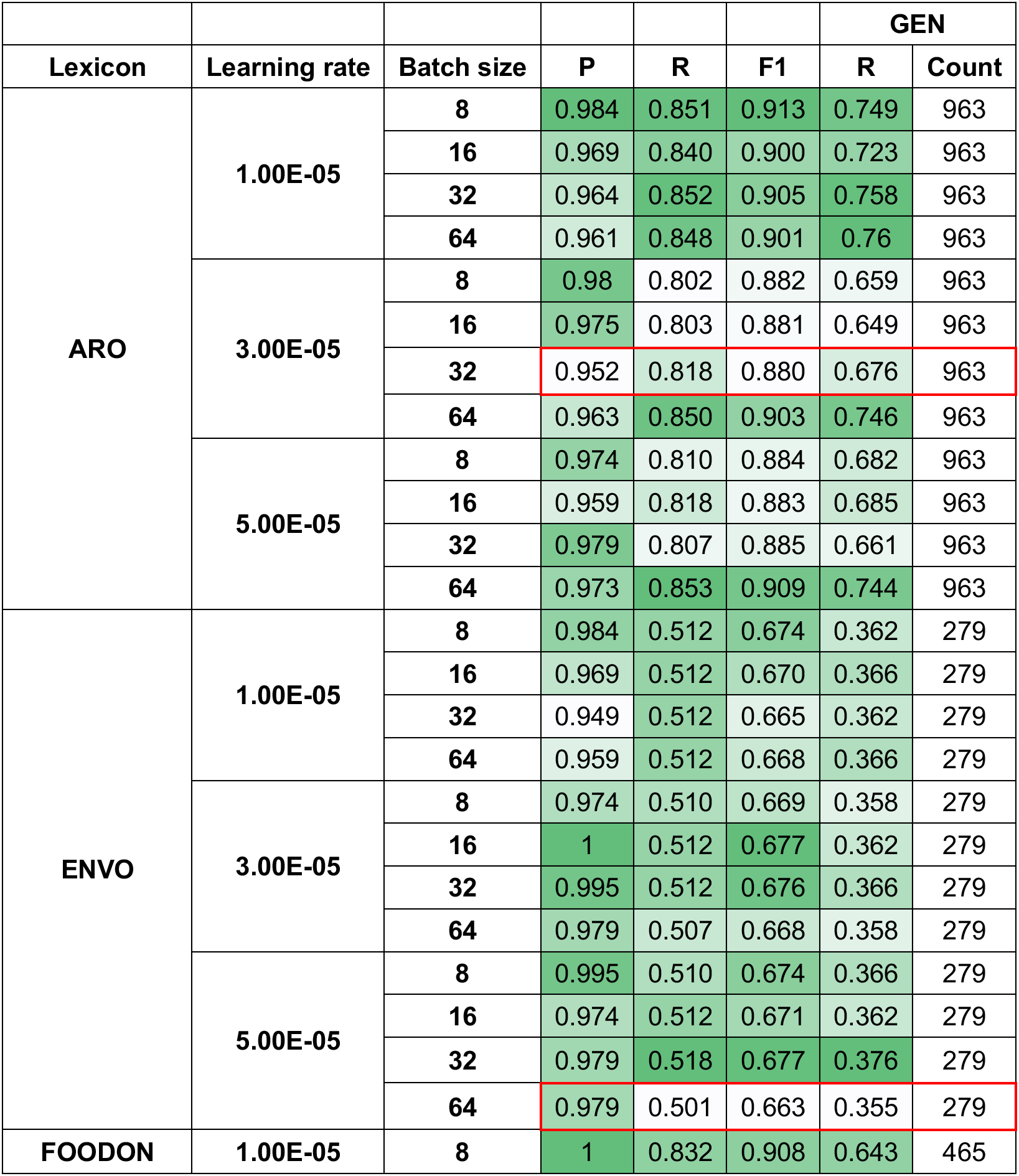
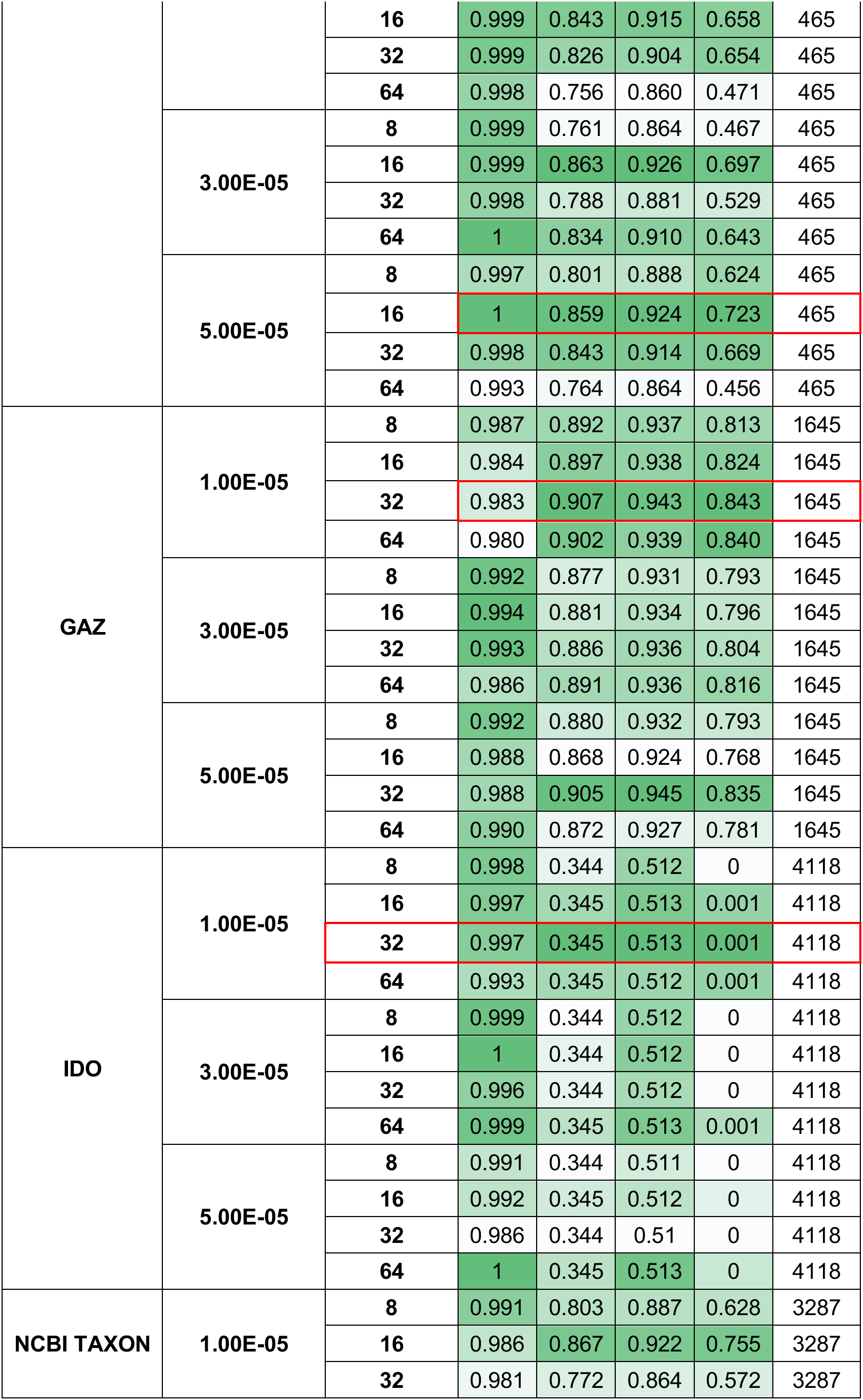
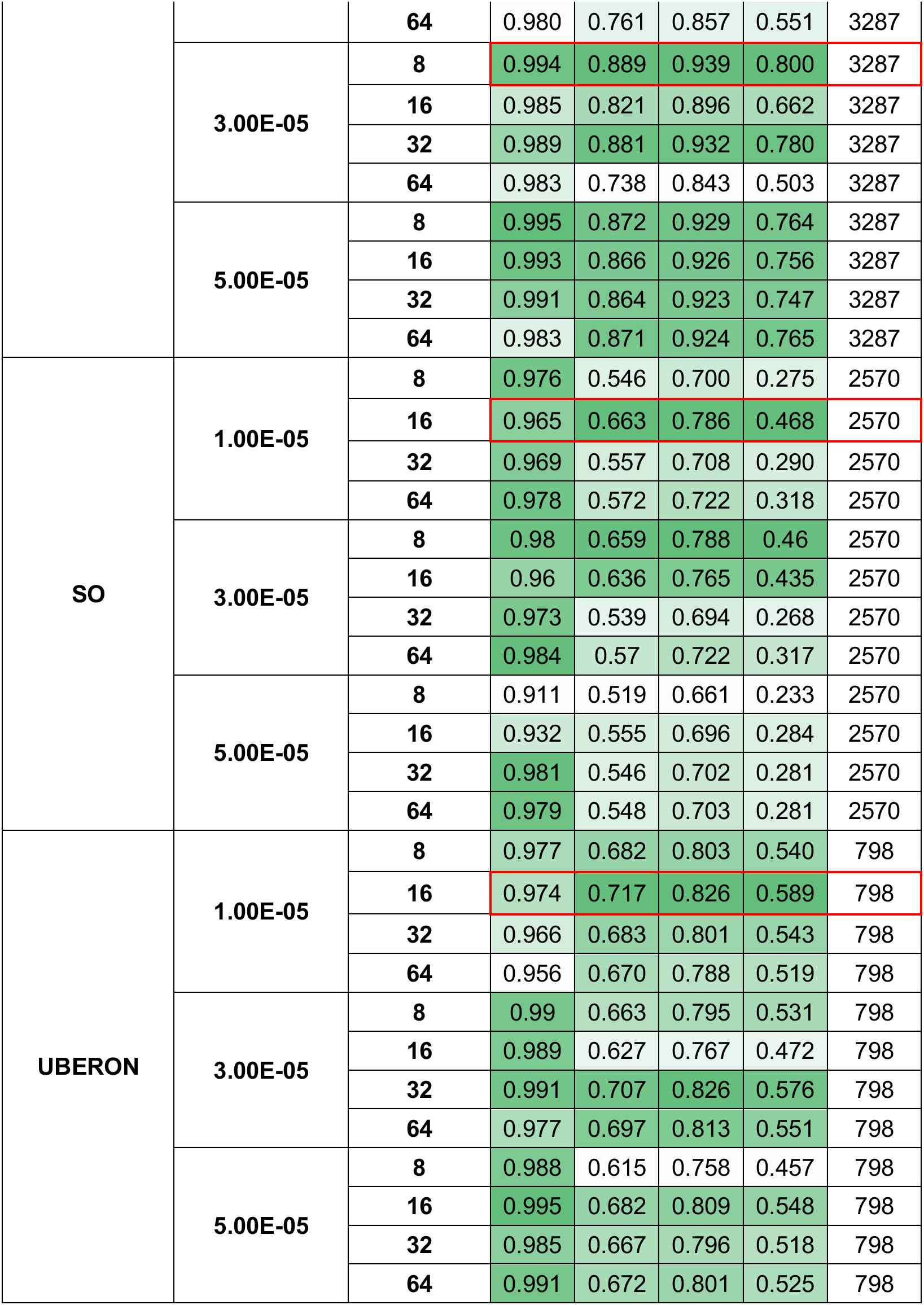
NER model performance and generalization ability. Precision (P), recall (R), and F1 scores were calculated on the testing NER dataset for the respective lexicon. Additionally, recall was measured on a subset of the training data: generalization (GEN). GEN counts were generated when neither the term nor their synonym appeared in the training set but appeared in the testing set. NER models were trained on a combination of three learning rates (1e-5, 3e-5, and 5e-5) and four batch sizes (8, 16, 32, 64). All models were trained using an epoch of 4. Boxed in red are the best-performing models based on generalizability and overall performance used for downstream NER annotations of PubMed. Green color scales were applied column-wise within lexicon boundaries to indicate the best-performing metric.

The 204,094 publications identified from the 2021 PubMed baseline by the logistic regression classifier were annotated using the NER models, generating over 2.2 million annotations (**Table 4**). These annotations were mapped to their corresponding ontology terms using a combination of Boolean and fuzzy-logic techniques, yielding 20,055 unique terms (**Figure S1**).

**Table 4:** NER predictions on 204k PubMed publication abstracts. Each lexicon’s best-performing NER model generated annotations on 204k PubMed abstracts. The total number of annotations, unique number of terms, and publications associated with each lexicon.

|  | **Annotations** | **Total terms** | **Publications** |
| --- | --- | --- | --- |
| **ARO** | 112,059 | 4,194 | 29,151 |
| **ENVO** | 78,298 | 188 | 40,654 |
| **FOODON** | 111,696 | 587 | 32,673 |
| **GAZ** | 223,487 | 5,672 | 80,475 |
| **IDO** | 400,711 | 197 | 133,420 |
| **NCBI TAXON** | 681,153 | 6,847 | 163,418 |
| **SO** | 461,144 | 678 | 105,236 |
| **UBERON** | 185,156 | 1,692 | 78,257 |
| **Total unique** | 2,253,704 | 20,055 | 201,791 |

Models for RE were evaluated based on loss, accuracy, and F1 score. Due to the relatively small size of some evaluation datasets (i.e., the smallest dataset, ENVO, with only 97 examples), accuracy and F1 scores exhibited limited variation across different learning rate and batch size combinations. The best-performing models were selected based on the highest accuracy and F1 scores, followed by the lowest loss (**Table 5**). The 2.2 million NER annotations were used to generate RE data, in which all ARO:other lexicon annotations found in the same sentence were masked and provided to the RE models for prediction, yielding 110,369 positive relationships across 33,580 relationship pairs (**Table 6**, **Figure 3**). These data, broken down by CARD’s drug classes, resistance mechanisms, and gene families are available in **Figure S2**. Frequencies by epidemiological term with geographical co-occurrence are available in **Tables S3-S7**.

**Figure 3.**
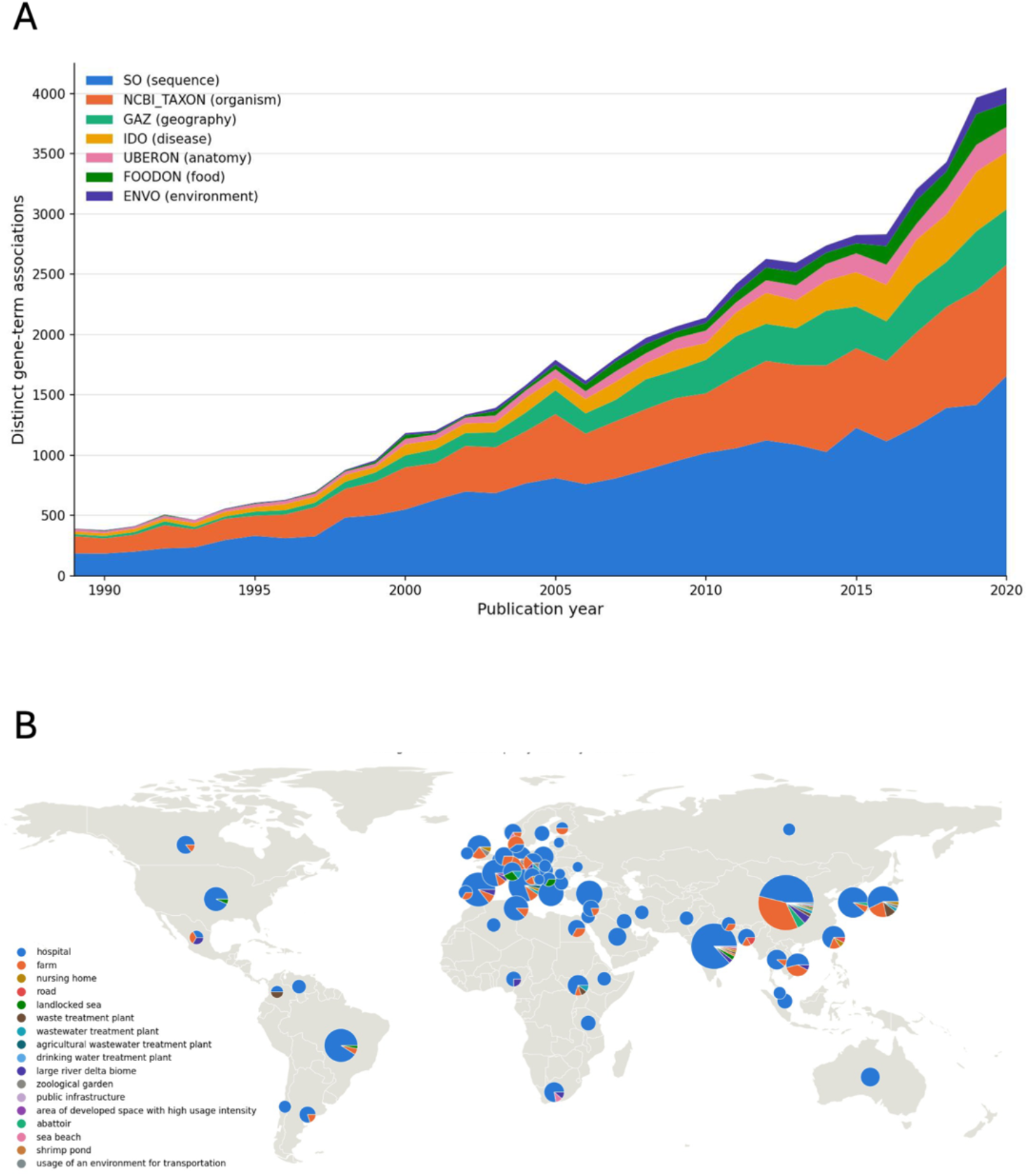
Relationship extraction (ARG:term) plotted by (A) publication date and lexicon, as well as (B) by country with pie charts representing most abundant ENVO terms. Figures created with the assistance of Anthropic Claude Sonnet version 4.6.

**Table 5:** RE model performance. Loss, accuracy, and F1 values were calculated based on predictions made on the evaluation dataset. The number of relationships found in each lexicon’s RE evaluation dataset is shown. RE models were trained on a combination of three learning rates (1e-5, 3e-5, and 5e-5) and four batch sizes (8, 16, 32, 64). All models were trained using an epoch of 4. Boxed in red are the best-performing models based on the lowest loss and overall performance used for downstream RE predictions. Green color scales were applied column-wise within lexicon boundaries to indicate the best-performing metric.

| Lexicon | Learning rate | Batch size | # Examples | Loss | Accuracy | F1 |
| --- | --- | --- | --- | --- | --- | --- |
| ENVO | 1.00E-05 | 8 | 97 | 0.289 | 0.897 | 0.945 |
|  |  | 16 |  | 0.345 | 0.897 | 0.945 |
|  |  | 32 |  | 0.347 | 0.897 | 0.945 |
|  |  | 64 |  | 0.403 | 0.897 | 0.945 |
|  | 3.00E-05 | 8 |  | 0.339 | 0.897 | 0.943 |
|  |  | 16 |  | 0.334 | 0.897 | 0.945 |
|  |  | 32 |  | 0.368 | 0.897 | 0.945 |
|  |  | 64 |  | 0.335 | 0.897 | 0.945 |
|  | 5.00E-05 | 8 |  | 0.475 | 0.914 | 0.952 |
|  |  | 16 |  | 0.334 | 0.897 | 0.943 |
|  |  | 32 |  | 0.338 | 0.897 | 0.945 |
|  |  | 64 |  | 0.353 | 0.897 | 0.945 |
| FOODON | 1.00E-05 | 8 | 110 | 0.284 | 0.909 | 0.952 |
|  |  | 16 |  | 0.275 | 0.909 | 0.952 |
|  |  | 32 |  | 0.295 | 0.909 | 0.952 |
|  |  | 64 |  | 0.336 | 0.909 | 0.952 |
|  | 3.00E-05 | 8 |  | 0.342 | 0.894 | 0.942 |
|  |  | 16 |  | 0.284 | 0.864 | 0.926 |
|  |  | 32 |  | 0.22 | 0.909 | 0.952 |
|  |  | 64 |  | 0.198 | 0.909 | 0.952 |
|  | 5.00E-05 | 8 |  | 0.421 | 0.894 | 0.942 |
|  |  | 16 |  | 0.295 | 0.879 | 0.932 |
|  |  | 32 |  | 0.256 | 0.864 | 0.926 |
|  |  | 64 |  | 0.188 | 0.909 | 0.952 |
| GAZ | 1.00E-05 | 8 | 198 | 0.538 | 0.866 | 0.928 |
|  |  | 16 |  | 0.408 | 0.874 | 0.933 |
|  |  | 32 |  | 0.392 | 0.874 | 0.933 |
|  |  | 64 |  | 0.369 | 0.874 | 0.933 |
|  | 3.00E-05 | 8 |  | 0.546 | 0.874 | 0.932 |
|  |  | 16 |  | 0.452 | 0.866 | 0.927 |
|  |  | 32 |  | 0.388 | 0.866 | 0.928 |
|  |  | 64 |  | 0.393 | 0.874 | 0.933 |
|  | 5.00E-05 | 8 |  | 0.554 | 0.874 | 0.932 |
|  |  | 16 |  | 0.479 | 0.874 | 0.932 |
|  |  | 32 |  | 0.384 | 0.891 | 0.94 |
|  |  | 64 |  | 0.39 | 0.866 | 0.928 |
| IDO | 1.00E-05 | 8 | 365 | 0.408 | 0.872 | 0.926 |
|  |  | 16 |  | 0.415 | 0.845 | 0.912 |
|  |  | 32 |  | 0.434 | 0.831 | 0.906 |
|  |  | 64 |  | 0.497 | 0.817 | 0.899 |
|  | 3.00E-05 | 8 |  | 0.520 | 0.904 | 0.943 |
|  |  | 16 |  | 0.398 | 0.904 | 0.943 |
|  |  | 32 |  | 0.329 | 0.881 | 0.93 |
|  |  | 64 |  | 0.459 | 0.845 | 0.911 |
|  | 5.00E-05 | 8 |  | 0.449 | 0.900 | 0.94 |
|  |  | 16 |  | 0.545 | 0.881 | 0.931 |
|  |  | 32 |  | 0.405 | 0.872 | 0.924 |
|  |  | 64 |  | 0.432 | 0.845 | 0.912 |
| NCBI TAXON | 1.00E-05 | 8 | 837 | 0.326 | 0.912 | 0.947 |
|  |  | 16 |  | 0.262 | 0.914 | 0.948 |
|  |  | 32 |  | 0.298 | 0.896 | 0.939 |
|  |  | 64 |  | 0.327 | 0.875 | 0.928 |
|  | 3.00E-05 | 8 |  | 0.274 | 0.936 | 0.961 |
|  |  | 16 |  | 0.231 | 0.94 | 0.964 |
|  |  | 32 |  | 0.271 | 0.912 | 0.947 |
|  |  | 64 |  | 0.293 | 0.886 | 0.932 |
|  | 5.00E-05 | 8 |  | 0.275 | 0.946 | 0.967 |
|  |  | 16 |  | 0.213 | 0.944 | 0.966 |
|  |  | 32 |  | 0.282 | 0.924 | 0.955 |
|  |  | 64 |  | 0.23 | 0.920 | 0.952 |
| SO | 1.00E-05 | 8 | 1,441 | 0.305 | 0.921 | 0.950 |
|  |  | 16 |  | 0.253 | 0.919 | 0.949 |
|  |  | 32 |  | 0.242 | 0.906 | 0.942 |
|  |  | 64 |  | 0.265 | 0.908 | 0.942 |
|  | 3.00E-05 | 8 |  | 0.350 | 0.927 | 0.954 |
|  |  | 16 |  | 0.297 | 0.925 | 0.952 |
|  |  | 32 |  | 0.22 | 0.929 | 0.955 |
|  |  | 64 |  | 0.217 | 0.926 | 0.953 |
|  | **5.00E-05** | **8** |  | 0.331 | 0.933 | 0.957 |
|  |  | **16** |  | 0.334 | 0.921 | 0.95 |
|  |  | **32** |  | 0.239 | 0.935 | 0.959 |
|  |  | **64** |  | 0.247 | 0.928 | 0.954 |
| **UBERON** | **1.00E-05** | **8** | **100** | 0.322 | 0.85 | 0.919 |
|  |  | **16** |  | 0.33 | 0.883 | 0.938 |
|  |  | **32** |  | 0.354 | 0.883 | 0.938 |
|  |  | **64** |  | 0.403 | 0.883 | 0.938 |
|  | **3.00E-05** | **8** |  | 0.487 | 0.833 | 0.902 |
|  |  | **16** |  | 0.33 | 0.833 | 0.907 |
|  |  | **32** |  | 0.335 | 0.867 | 0.929 |
|  |  | **64** |  | 0.346 | 0.883 | 0.938 |
|  | **5.00E-05** | **8** |  | 0.509 | 0.85 | 0.911 |
|  |  | **16** |  | 0.456 | 0.817 | 0.893 |
|  |  | **32** |  | 0.305 | 0.833 | 0.907 |
|  |  | **64** |  | 0.339 | 0.883 | 0.938 |

**Table 6.** Number of ARO-lexicon relationships generated from 204k publication abstracts. The number of unique relationship term pairs, ARO terms, total relationships, and publications associated with each lexicon is shown. In total, there were 33,580 unique pairs of ARO:lexicon terms and 110,369 relationships from 24,235 unique publications. Relationships that appeared multiple times in a single publication were counted as one relationship.

| **Lexicon** | **Relationship pairs** | **Relationships** | **Publications** | **Unique Epi terms** | **Unique ARO terms** |
| --- | --- | --- | --- | --- | --- |
| **ENVO** | 793 | 2,768 | 1,819 | 48 | 390 |
| **FOODON** | 1,185 | 2,742 | 1,255 | 93 | 373 |
| **GAZ** | 4,233 | 7,706 | 4,000 | 650 | 775 |
| **IDO** | 2,829 | 9,931 | 5,620 | 45 | 1,148 |
| **NCBI TAXON** | 8,454 | 32,833 | 15,090 | 883 | 2,232 |
| **SO** | 14,240 | 50,338 | 16,171 | 253 | 2,960 |
| **UBERON** | 1,846 | 4,051 | 2,583 | 175 | 692 |
| **Total** | 33,580 | 110,369 | 24,235 | 2,147 | 3,528 |

### Data visualization

Having identified associations between ARGs and epidemiology terms from scientific literature (NER and RE data), we can leverage these associations to examine metrics for modelling ARG transmission based on the 2021 PubMed baseline. This enables a quantitative assessment of the relatedness between two terms, such as the similarity in ARG burden between “wastewater” and “hospital”. To quantify these similarities, we used two metrics: gene overlap and Jaccard similarity (**Table 7**). To visualize ARG transmission patterns using a Confusogram, where nodes represent different epidemiology terms and edges represent numbers of shared ARGs, we assessed publications that mentioned a specific term of interest (e.g., *E. coli*) and gathered all those PubMed identifiers (PMIDs) and their subsequent annotations and relationships. **Figure 4** displays a Confusogram capturing the highest Jaccard similarity scores among publications that mention *E. coli*, illustrating an abundance of terms from FOODON, indicating that ARGs are likely being transmitted among food and animals. Furthermore, a UMAP visualization pipeline was designed to analyze transmission pathways among entire ontologies, in which the plot displays the clustering of epidemiological terms based on shared ARGs (**Figure 5**). If more terms cluster together, it indicates that they share more ARGs based on the RE data, e.g., seafood terms like “tilapia”, “mussel”, and “prepared seafood product” cluster together.

**Table 7.** Example similarity metrics among relationship terms. Gene overlap counts the number of ARGs shared among two terms (e.g., “*Escherichia coli*” and “Canada” share 8 ARGs in abstracts that have the word “Canada” in them). Jaccard similarity assesses the gene overlap and the total number of ARGs associated with both terms.

| Lexicon-Lexicon | Term 1 | Term 2 | Gene Overlap Count | Jaccard Similarity |
| --- | --- | --- | --- | --- |
| NCBI_TAXON-GAZ | *Escherichia coli* | Canada | 8 | 0.3478 |
| NCBI_TAXON-GAZ | *Enterococcus faecium* | United States | 1 | 0.25 |
| NCBI_TAXON-GAZ | *Salmonella* | United States | 1 | 0.25 |
| NCBI_TAXON-GAZ | *Salmonella* | Toronto | 1 | 0.25 |
| GAZ-ENVO | Toronto | hospital | 1 | 0.25 |
| GAZ-ENVO | India | hospital | 1 | 0.25 |
| GAZ-ENVO | United States | hospital | 1 | 0.25 |
| GAZ-GAZ | United States | Toronto | 1 | 0.25 |
| NCBI_TAXON-ENVO | *Salmonella* | hospital | 1 | 0.25 |
| NCBI_TAXON-GAZ | *Salmonella* | Province of Ontario | 1 | 0.20 |

**Figure 4.**
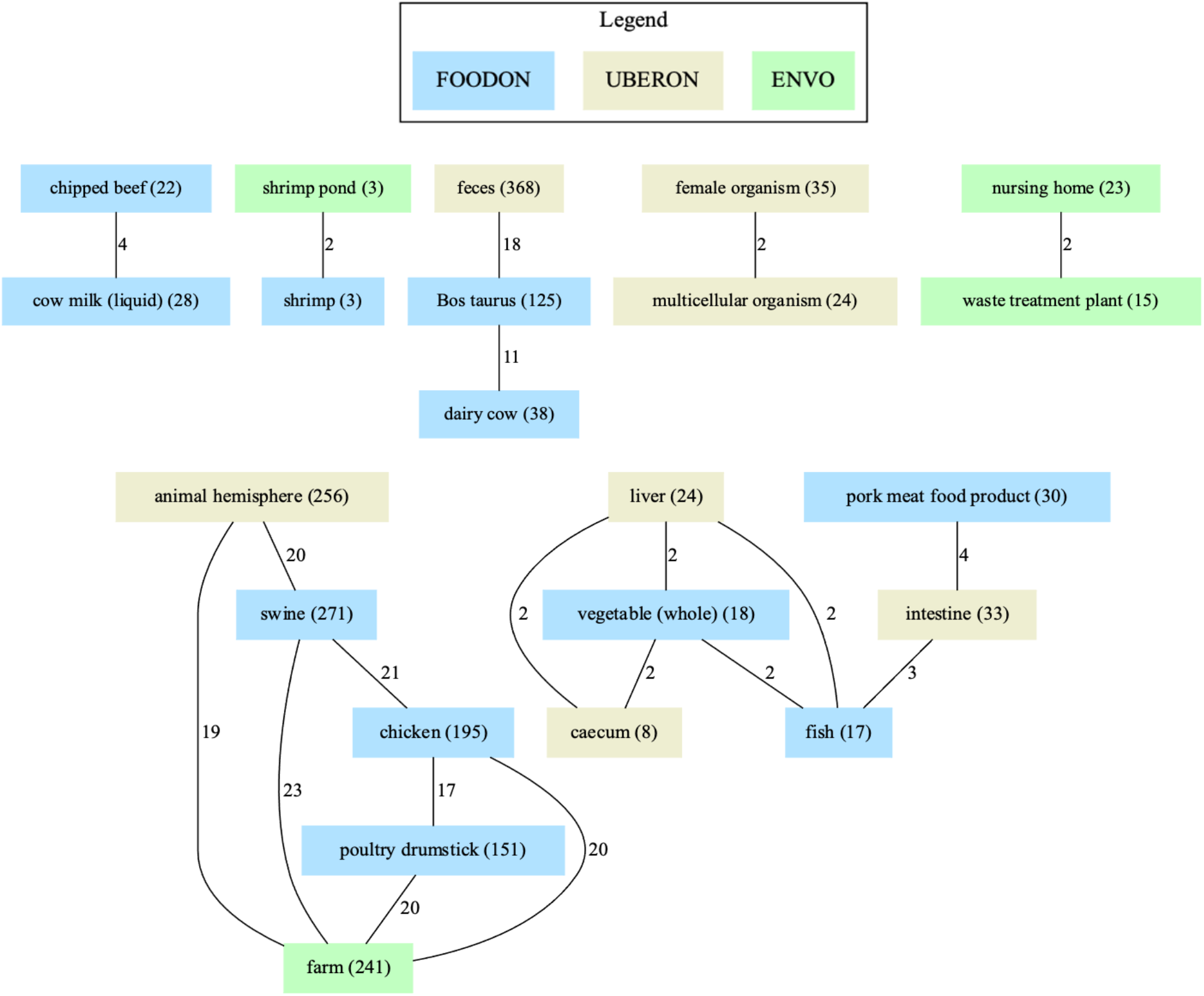
*Escherichia coli* confusogram using the abstract dataset’s uber-anatomy (UBERON), environment (ENVO), and food ontologies (FOODON). Each edge indicates the number of shared ARGs (from the ARO) between two epidemiological terms. Each node includes the term name and the number of publications with the respective term.

**Figure 5.**
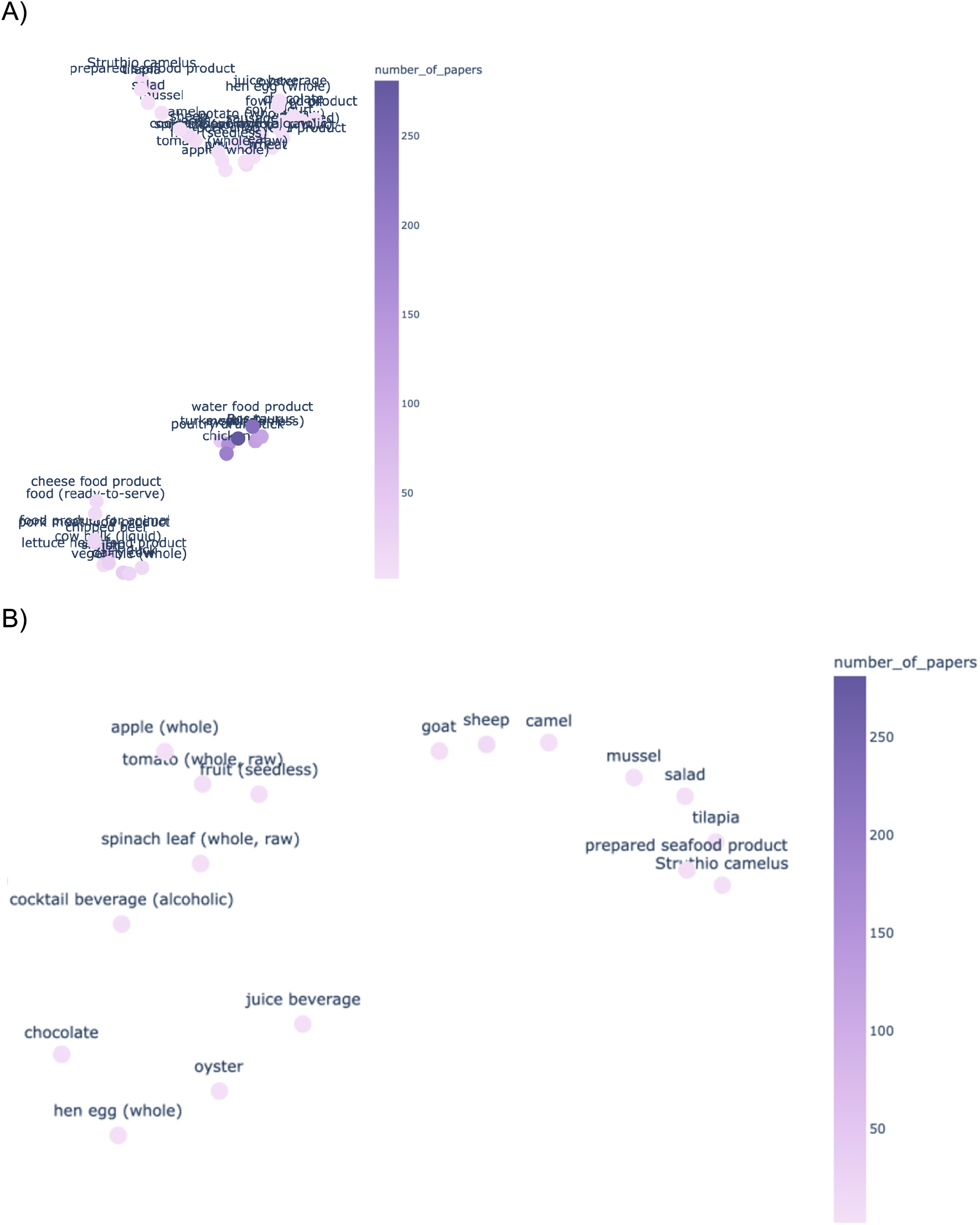
UMAP visualization for FOODON terms found in 204k PubMed abstracts. The lighter the dot, the fewer publications that mention the term. A) entire visual of all FOODON terms, illustrating three distinct clusters, with B) a re-oriented close-up of the upper cluster.

### Integration with the Comprehensive Antibiotic Resistance Database

All data are available for download at the CARD website (see Software & Data Availability). In addition, the RE results have been incorporated into CARD’s Broad Street data schema so web pages for individual ARGs are now accompanied by CARD:Epi data (**Figure 6**).

**Figure 6.**
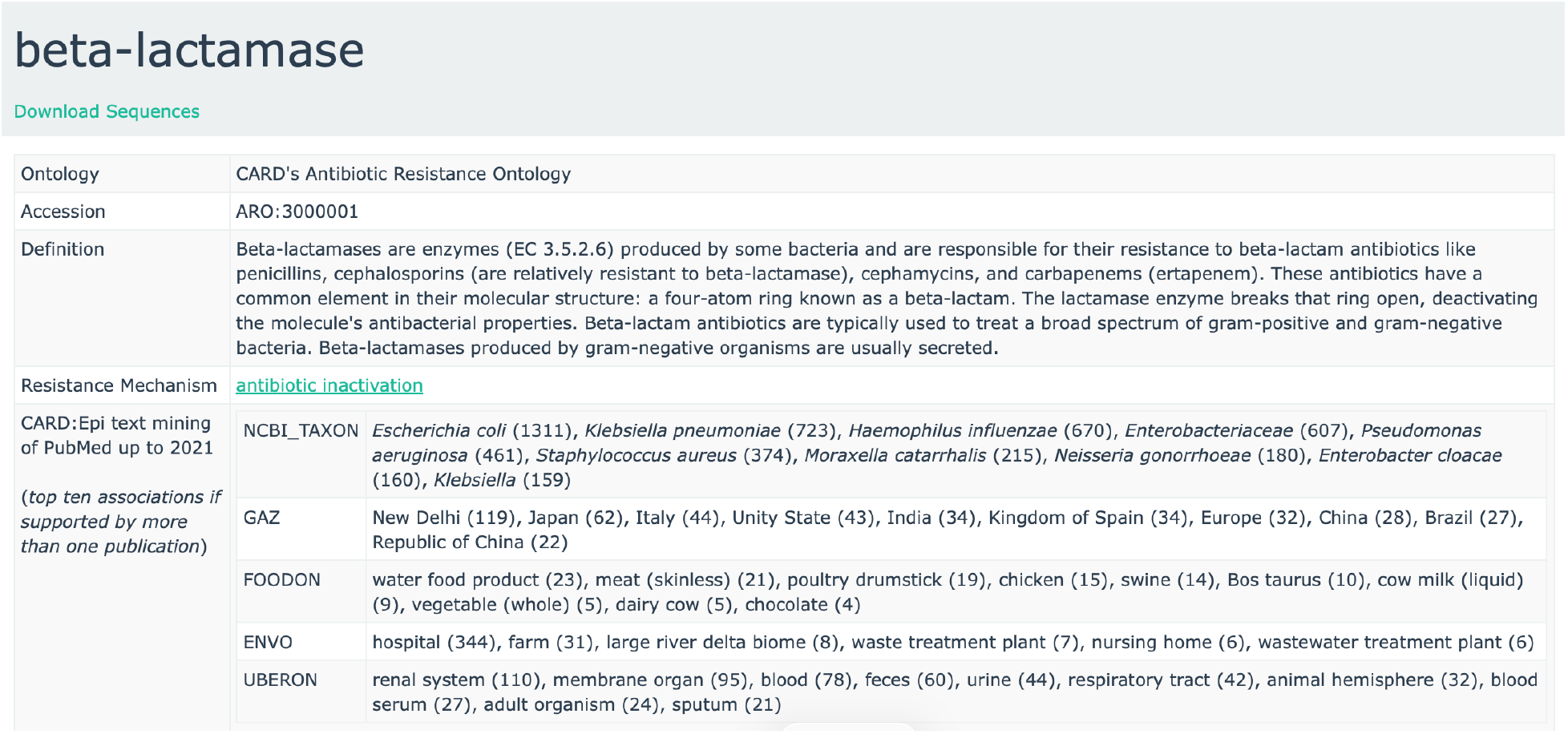
Incorporation of the CARD:Epi data at the CARD website, showing results for ARO:3000001 “beta-lactamase”.

## DISCUSSION

While genomic surveillance and molecular epidemiology of AMR determinants is increasingly routine, this simply increases the need for contextual data for interpretation of results, particularly for the gene lists generated by genomic and metagenomic studies. While CARD’s canonical data (gene sequences and mutations, plus their association with antibiotics) is hand-curated from the scientific literature to ensure high-quality reference data, similar curation of the total AMR literature to mine epidemiological associations is not practical. Building upon past efforts using machine learning to triage literature for human curators, i.e., CARD*Shark^32^, CARD:Epi reflects CARD’s first deployment of NLP to generate high-quality contextual data sets, with multiple steps of human evaluation to ensure trustworthy results. As biocurators, our focus has been on the synthesis of high-quality contextual data for AMR genomics and their incorporation into CARD, with planned annual updates. We look forward to their incorporation by us and others into AMR bioinformatics tools and modelling efforts.

To generate NER and subsequently RE training datasets, we used standard sets of terms that represent concepts that can be identified within the biomedical literature. This was necessary when normalizing predictions from NER models back to a standard set of terms. We found two major factors that impacted how functional these ontologies were for text mining purposes: (i) how polysemantic the ontology terms were, and (ii) how comprehensive the ontology was and the depth/nuance of terms within.

Pattern-matching algorithms were used in building gold-standard training datasets, however they did not consider that a term may have multiple meanings, thus many annotations resulted in a mismatch between an annotated term and the corresponding lexicon term’s meaning. For example, the term “an” in FOODON refers to a sweet bean paste, but this is not the most common meaning in biomedical text (which is as an indefinite article used before singular nouns), resulting in mismatching. The ontologies most impacted by a mismatch were GAZ, UBERON, and SO. Given the epidemiological importance of geography, we are exploring more detailed alternatives to GAZ. Ontologies with terms that have only one definition were mismatched much less often and this was seen for ARO, NCBI TAXON, and IDO, which lost very few annotations to filtering.

While polysemantic terms frequently cause mismatches, an inability to identify relevant terms within a text was another concern. There were three reasons we could not identify terms within text: (i) the ontology was not comprehensive enough and lacked relevant terms, (ii) the ontology terms were too verbose, or (iii) the ontology did not capture enough nuance within the terms. After filtering, many terms were manually added to ENVO, FOODON, and NCBI TAXON, indicating that the ontologies alone could not capture the extent of terms belonging to their overall concept. For NCBI TAXON, the ontology was not comprehensive and lacked species abbreviations. These abbreviations were added retroactively to capture all bacterial taxonomy. While comprehensive, FOODON had verbose terms and went into too much detail to be helpful for a text-mining application. For example, the term “poultry” is not in FOODON, but the terms “poultry product”, “poultry (frozen)”, “poultry (raw)” and many other verbose terms for poultry food items can be found in the ontology. As NLP benefits from both specific and general terms, curators should evaluate their reference ontologies prior to use.

Both mismatches and the lack of matches stem from an ontology’s structure and the terms contained within. These annotation problems were corrected via manual filtering steps (the removal of terms) as well as identifying missing terms that require one to review annotations sentence-by-sentence to improve the resulting NER training/testing datasets. Creating gold-standard datasets is a manual process, which is time-consuming and requires attention to detail and patience. We have generated gold-standard datasets that identify epidemiology concepts alongside ARGs and the relationships between them. Other NLP researchers can use these data to train new models and benchmark current models to understand how these terms interact in biomedical literature, among many other applications.

The way ontologies are structured also has an impact on downstream tasks like visual representation of ARG transmission patterns (via a Confusogram or UMAP). The goal of these visuals is to draw conclusions between ARGs and epidemiological data (i.e., environments, food/animals, etc., geographical locations, etc.) found in biomedical literature. Generating a visual that encapsulates a specific term (e.g., ARG transmission patterns for the term “*Escherichia coli*”) can be simple. However broader concepts, e.g., ARG transmission patterns for a country, may pose a challenge due to the lack of harmonization within the data. For example, when assessing ARG transmission patterns in Canada, the dataset separately contains “Canadian”, “Toronto”, “Province of Ontario” and other terms that all resolve to Canada. Finding an efficient way to harmonize a domain-specific concept across a variety of lexicons poses a challenge, perhaps solvable by next generation large language models (LLMs).

Our datasets hold a large amount of data from 204k PubMed abstracts, however when we explore more specific epidemiological hypotheses of ARG transmission (e.g., predicting ARG transmission through seafood importation routes), there is a lack of relationships among epidemiology terms and ARGs (**Figure S3**). The thinness of these data reflects bias in AMR research towards clinical settings (**Figure 3)**, BioBERT’s dependence on finding relationships within individual sentences but not among sentences, and our restriction to analyzing abstracts, e.g., our analysis found 26 publications supporting association of ARGs with shrimp, but only 3 with tilapia. In the future, we will turn to extracting information from PubMed Central full-text articles to sample more relationships to answer these questions.

## CONCLUSIONS

We conducted an interdisciplinary analysis of the scientific literature using eight ontologies to understand AMR epidemiology, generating gold-standard datasets and training BioBERT NER and RE models to extract information regarding AMR determinants and their epidemiology. This dataset can be manipulated in many ways to potentially aid in modelling ARG transmission pathways among environments, locations, years, foods, and animals. To represent these data, we have designed two visualization methods via a Confusogram and UMAP to analyze and predict pathways of AMR determinants found within the CARD:Epi dataset. Overall, this work has indicated that a copious amount of untapped epidemiological data exists in biomedical text that can be mined by leveraging NLP.

## SOFTWARE & DATA AVAILABILITY

All training and testing data set files, the AMR-Epi classifier model files, NER and RE model files, and 2021 PubMed baseline results files can be downloaded from the Comprehensive Antibiotic Resistance Database (https://card.mcmaster.ca/download). Python scripts for visualization of results can be found in GitHub repository https://github.com/tiffanyta1402/cardepi-analysis.

## SUPPLEMENTARY DATA

Supplementary Data are available online in association with this publication.

## ACKNOWLEDGEMENTS

Computational support was provided by the McMaster Faculty of Health Sciences Advanced Computing Facility, supplemented by hardware donations and loans from Cisco Systems Canada, Hewlett Packard Enterprise, and Pure Storage. Figures 3 and S2 were created with the assistance of Anthropic Claude Sonnet version 4.6. AGM currently holds McMaster’s David Braley Chair in Computational Biology, generously supported by the family of the late Mr. David Braley. Drs. John Nash, Guillaume Paré, and Lori Burrows provided constructive feedback on this research. Comments on this manuscript were provided by Brian P. Alcock.

## AUTHOR CONTRIBUTIONS

AE: conceptualization, methodology, software, validation, formal analysis, investigation, data curation, writing - original draft, visualization, writing - review & editing. TET: methodology, software, investigation, data curation, visualization, writing - original draft, writing - review & editing. CZ, AI, RU, SR: validation. AGM: conceptualization, resources, supervision, project administration, funding acquisition, writing - review & editing. AE and TET contributed equally to this work.

