## Supplementary Tables S1-2, Figures S1-S3 for "CARD:Epi – Contextualizing Antimicrobial Resistance Determinants Using Deep Learning Language Models"

Table S1: Breakdown of ontologies used, the concepts they represent, and the parent term used as primary filter.

| Lexicon | **Main concept** | **Parent filter** |
|---|---|---|
| ARO | ARG names | Determinant of antibiotic resistance (ARO:3000000) |
| FOODON | Food products | Food product (FOODON:00001002) |
| UBERON | Anatomy parts | Anatomical entity (UBERON:0001062) |
| ENVO | Environments | Construction (ENVO:01001813) |
| NCBI TAXON | Bacterial taxonomy | Bacteria (0) |
| GAZ | Geographical locations | No filter |
| SO | Genomic terms | No filter |
| IDO | Infectious disease terms | Continuant (BFO:0000002) |

Table S2: Relationship counts within the RE training and testing dataset. ARO to lexicon relationships were generated using sentences that contain at least one ARO term and another term from a different lexicon. The total number of ARO:lexicon relationships available can be seen in the "All" column. Relationships were manually labelled and given a 0/1 depending on whether the ARO:lexicon terms contextualize one another (1) or not (0). The subset of relationships labelled 1 was used to train the RE models.

| Lexicon | All | Labelled 0 | Labelled 1 | Unique Publications |
|---|---|---|---|---|
| **ENVO** | 386 | 25 | 361 | 153 |
| **FOODON** | 438 | 49 | 389 | 142 |
| **GAZ** | 791 | 62 | 729 | 339 |
| **IDO** | 1,460 | 273 | 1,187 | 549 |
| **NCBI TAXON** | 3,347 | 583 | 2,763 | 968 |
| **SO** | 5,766 | 1,204 | 4,558 | 1,149 |
| **UBERON** | 400 | 58 | 342 | 203 |
| **Total** | 12,588 | 2,254 | 10,329 | 1,664 |

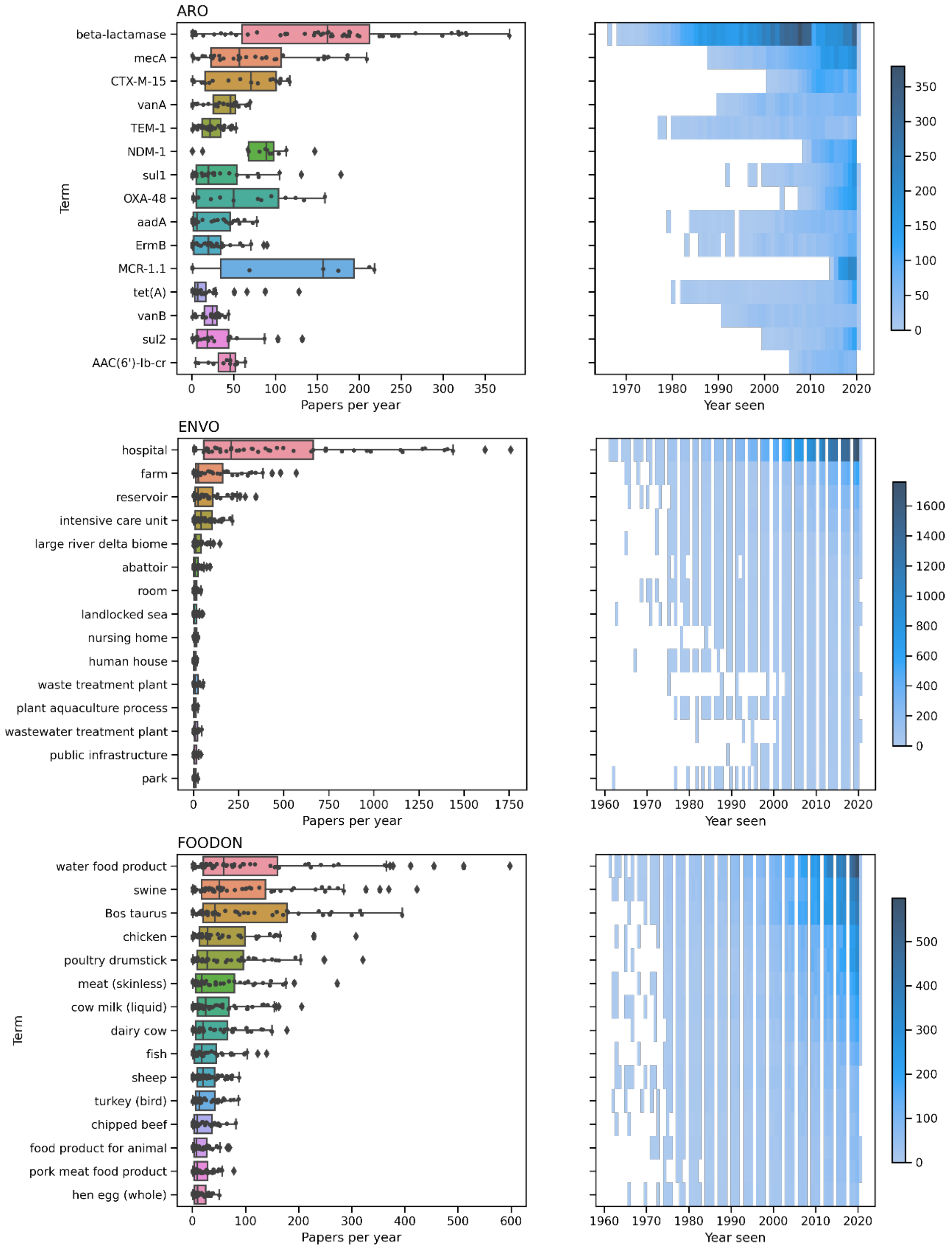

ARO
beta-lactamase
mecA
CTX-M-15
vanA
TEM-1
NDM-1
sul1
OXA-48
aadA
ErmB
MCR-1.1
tet(A)
vanB
sul2
AAC(6')-Ib-cr
Term
0 50 100 150 200 250 300 350
Papers per year
1970 1980 1990 2000 2010 2020
Year seen
0 50 100 150 200 250 300 350
ENVO
hospital
farm
reservoir
intensive care unit
large river delta biome
abattoir
room
landlocked sea
nursing home
human house
waste treatment plant
plant aquaculture process
wastewater treatment plant
public infrastructure
park
Term
0 250 500 750 1000 1250 1500 1750
Papers per year
1960 1970 1980 1990 2000 2010 2020
Year seen
0 200 400 600 800 1000 1200 1400 1600
FOODON
water food product
swine
Bos taurus
chicken
poultry drumstick
meat (skinless)
cow milk (liquid)
dairy cow
fish
sheep
turkey (bird)
chipped beef
food product for animal
pork meat food product
hen egg (whole)
Term
0 100 200 300 400 500 600
Papers per year
1960 1970 1980 1990 2000 2010 2020
Year seen
0 100 200 300 400 500

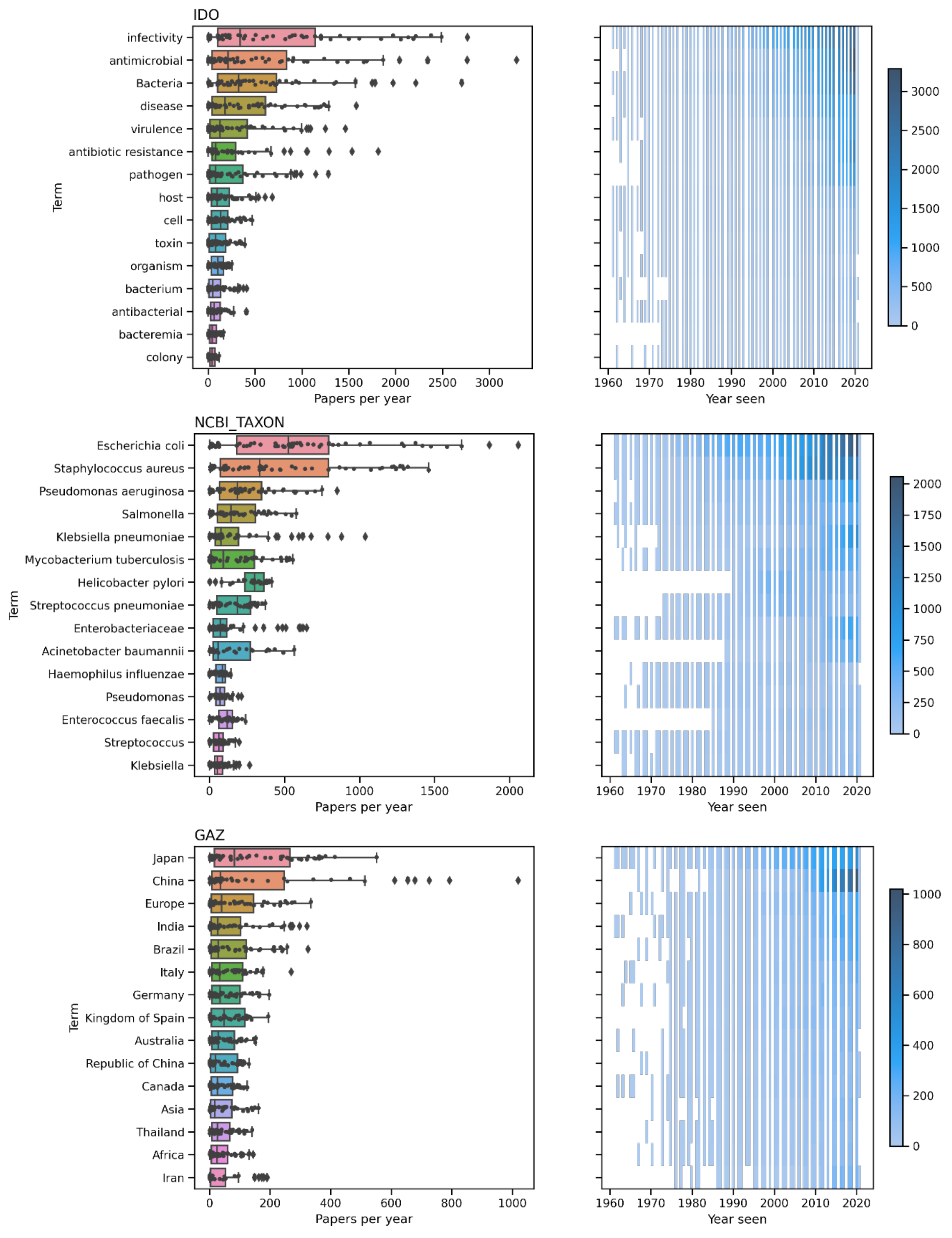
IDO
infectivity
antimicrobial
Bacteria
disease
virulence
antibiotic resistance
pathogen
host
cell
toxin
organism
bacterium
antibacterial
bacteremia
colony
Term
0
500
1000
1500
2000
2500
3000
Papers per year
1960
1970
1980
1990
2000
2010
2020
Year seen
0
500
1000
1500
2000
2500
3000
NCBI_TAXON
Escherichia coli
Staphylococcus aureus
Pseudomonas aeruginosa
Salmonella
Klebsiella pneumoniae
Mycobacterium tuberculosis
Helicobacter pylori
Streptococcus pneumoniae
Enterobacteriaceae
Acinetobacter baumannii
Haemophilus influenzae
Pseudomonas
Enterococcus faecalis
Streptococcus
Klebsiella
Term
0
500
1000
1500
2000
Papers per year
1960
1970
1980
1990
2000
2010
2020
Year seen
0
250
500
750
1000
1250
1500
1750
2000
GAZ
Japan
China
Europe
India
Brazil
Italy
Germany
Kingdom of Spain
Australia
Republic of China
Canada
Asia
Thailand
Africa
Iran
Term
0
200
400
600
800
1000
Papers per year
1960
1970
1980
1990
2000
2010
2020
Year seen
0
200
400
600
800
1000

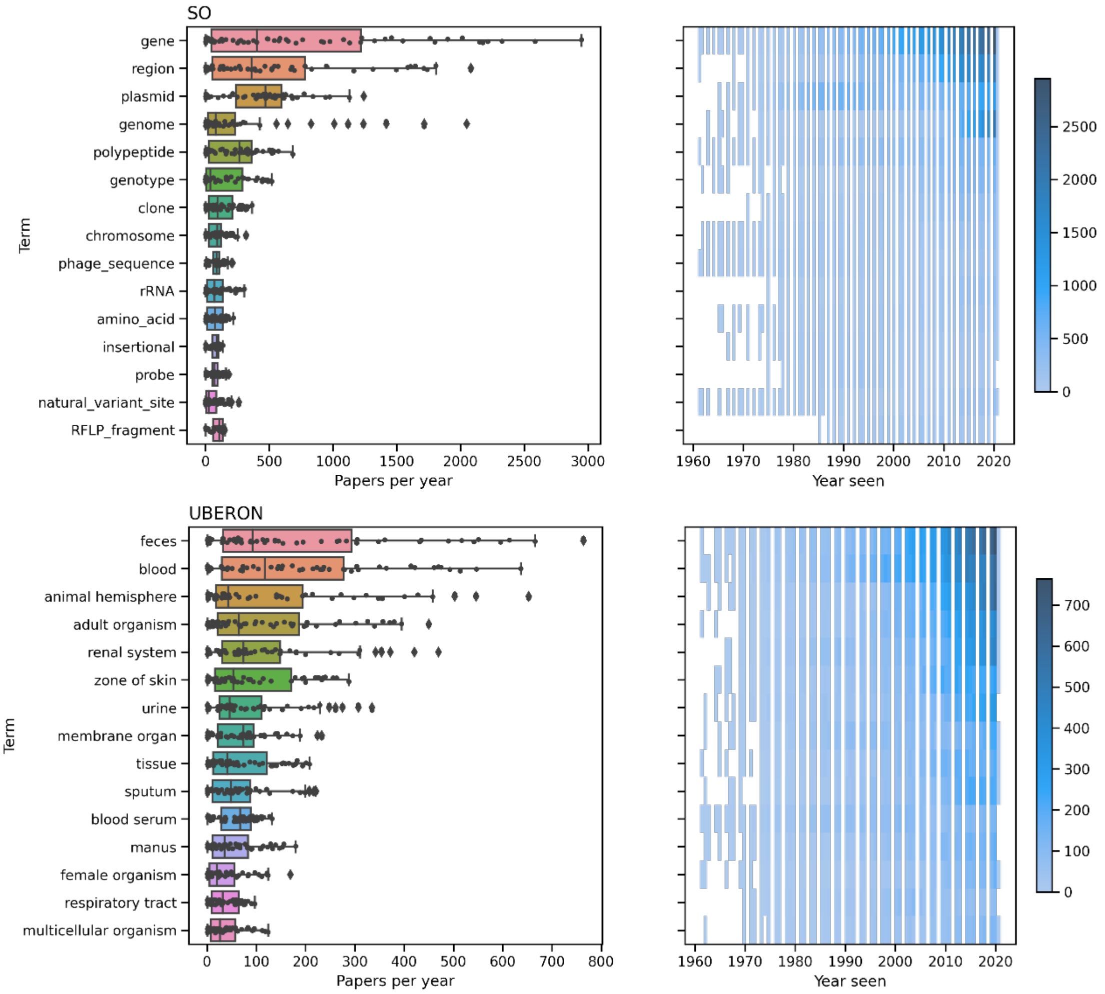


Figure S1: Top 15 terms in each lexicon annotated from 204k publications. A boxplot and heatmap represent the number of publications each term appears in per year, sorted in descending order by the total number of annotations.

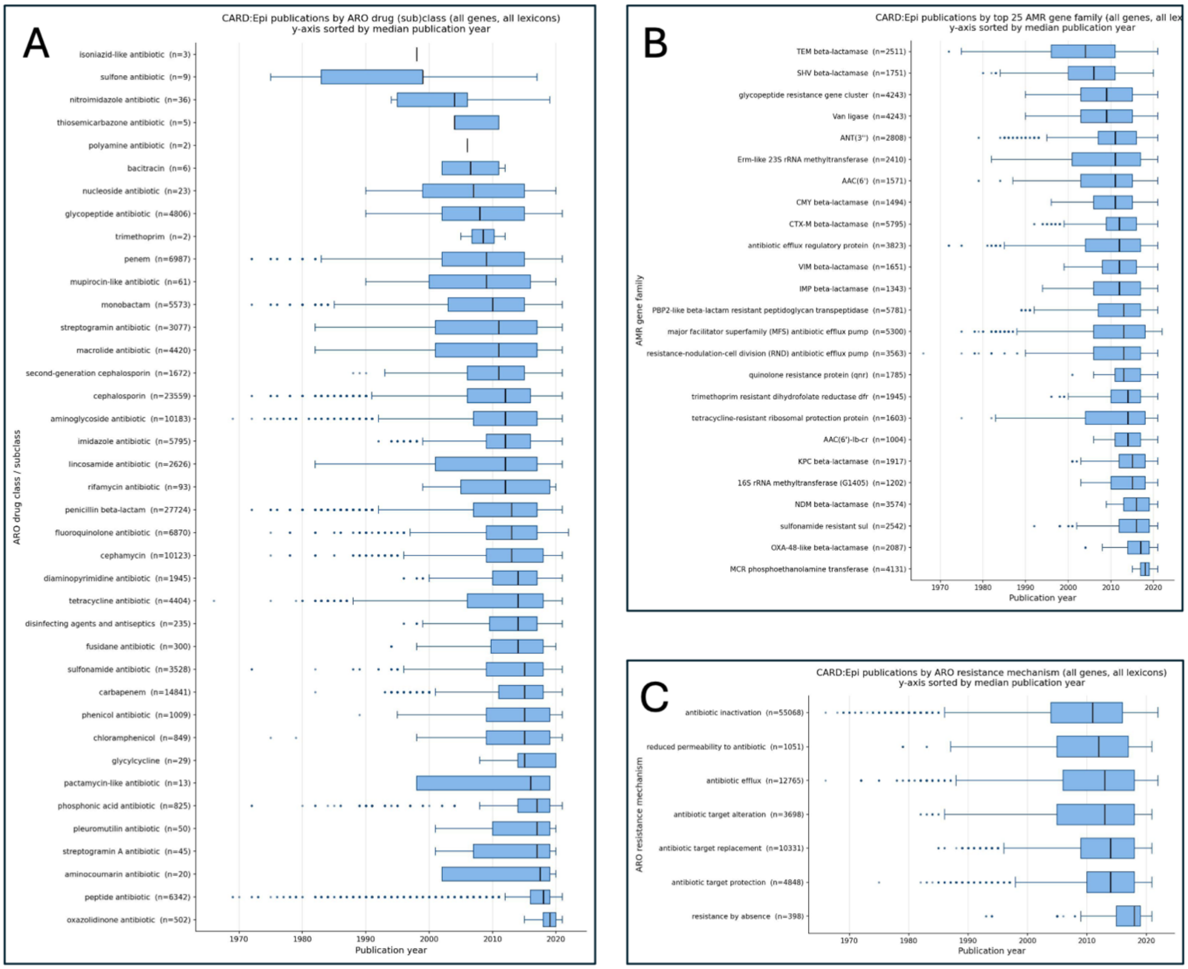


Figure S2: Box plots of publications by year for CARD drug classes (A), top 25 AMR gene families (B), and resistance mechanisms (C). Boxes span the 25th to 75th percentile, with the median indicated by a bold line. Whiskers represent the range of earliest and latest publication (1.5 interquartile range), with points representing outliers. Figure created with the assistance of Anthropic Claude Sonnet version 4.6.

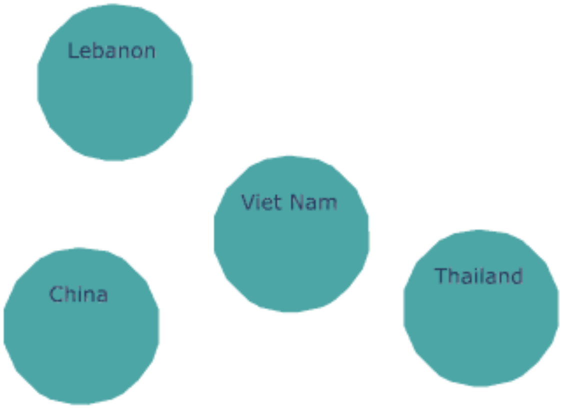


Figure S3: UMAP representation of seafood importation routes based on shared ARGs. A sub-dataset was created for all abstracts that contain seafood terms (e.g., fish, mussel, shrimp), then filtered to obtain all GAZ terms. Each term in this visual was mentioned in only a single abstract.
